# Exploring the only known case of sympatry in sportive lemurs: isolation by distance or speciation?

**DOI:** 10.64898/2026.08.27.747501

**Authors:** Jordi Salmona, Besoa Ranjavao, Emmanuel Rasolondraibe, Ando Nantenaina Rakotonanahary, Tantely Ralantoharijaona, Fabien Jan, Barbara Le Pors, Helena Teixeira, Célia Kun-Rodrigues, Mohamed Thani Ibouroi, Said Ali Ousseni Durham, Radavison Zaranaina, Vivien Gabillaud, Mélanie Barnavon, Angelika Beck, Ana Rita Monteiro, Isa Aleixo-Pais, Ana Priscilla Sousa, Paul Hohenlohe, Stéphanie M. Carrière, Solofo Rakotondraompiana, Tendro Radanielina, Sébastien Wohlhauser, Patrick Ranirison, Nicole Volasoa Andriaholinirina, Romule Rakotondravony, Solofonirina Rasoloharijaona, Rasmus Heller, John Rigobert Zaonarivelo, Gabriele Maria Sgarlata, Lounès Chikhi

**Author notes:** JS and BR should be considered joint first author. corresponding authors: JS.

## Abstract

Among Madagascar primates, the sportive lemurs (family Lepilemuridae) have seen their species diversity increase from eight in 2005 to 26 in 2009 mostly by applying the phylogenetic species concept to DNA barcode data. Despite the genus being speciose, only one case of sympatry is known from northern Madagascar, where two sportive lemur species described based on low mtDNA divergence, *Lepilemur ankaranensis* and *Lepilemur milanoii*, were found to co-occur at the center of their joint distribution range. Here, to clarify the taxonomy of these two species and examine their sympatry, we apply an integrative taxonomic framework to genomic and morphological data from 84 individuals of *L. ankaranensis* and *L. milanoii*, encompassing their entire distribution range and the forest of Analafiana, beyond their southernmost limit. Using clustering, multivariate, and isolation by distance analyses, we find no evidence of a sympatric zone and show that despite clear genetic differentiation between regions, the genomic and morphological diversity of the *L. ankaranensis* - *L. milanoii* - Analafiana group is clinal and explained by geographic distance. These results clarify that *L. milanoii* is a junior synonym of *L. ankaranensis* and that the Analafiana forest population belongs to *L. ankaranensis*, extending its distribution. It further implies that the “sympatric” zone—the Andrafiamena forest—hosts conspecific individuals with slightly differentiated mtDNA backgrounds, rather than slightly differentiated sympatric species. Lastly, we re-evaluate the IUCN conservation metrics of *L. ankaranensis*, which continue to qualify as Endangered (EN) under the B1ab(i-v) criteria.

## Introduction

Madagascar is recognized as a major biodiversity hotspot (Myers et al. 2000), known for hosting an entire superfamily of highly diverse primates, the Lemuroidae. While 20 species of lemurs were recognized by primatologists in the 1980s (Tattersall, 1982), more than 100 species have been described (108 species, Mittermeier et al., 2023), following the introduction of mitochondrial DNA barcoding and extensive field efforts (e.g. Louis, 2006). This increase in the number of species has been particularly important in nocturnal lemurs such as mouse lemurs and sportive lemurs (genera *Microcebus* and *Lepilemur*, respectively). For instance, in sportive lemurs (family Lepilemuridae), a clade of medium-sized lemurs, the number of described species increased from 8 in 2005 to 26 in 2009 (Figure 1a, Table S1, Andriaholinirina et al., 2006; Craul et al., 2007; Louis, 2006), essentially by applying the phylogenetic species concept (PSC, Louis & Lei, 2014) to DNA barcode data. The PSC concept has been, however, criticized for overestimating species diversity, also referred to as taxonomic inflation (Tattersall, 2007, Markolf et al., 2011, van Elst et al., 2025). In a recent study, van Elst et al., (2025) have suggested that by applying an integrative approach linking molecular, morphological and spatial data, one could identify such an inflation in mouse lemurs, where the number of species increased from three to 25 between the 1990s and the 2020s. Their results suggested that the actual number of well-established species could be closer to 19 than 25.

**Figure 1:**
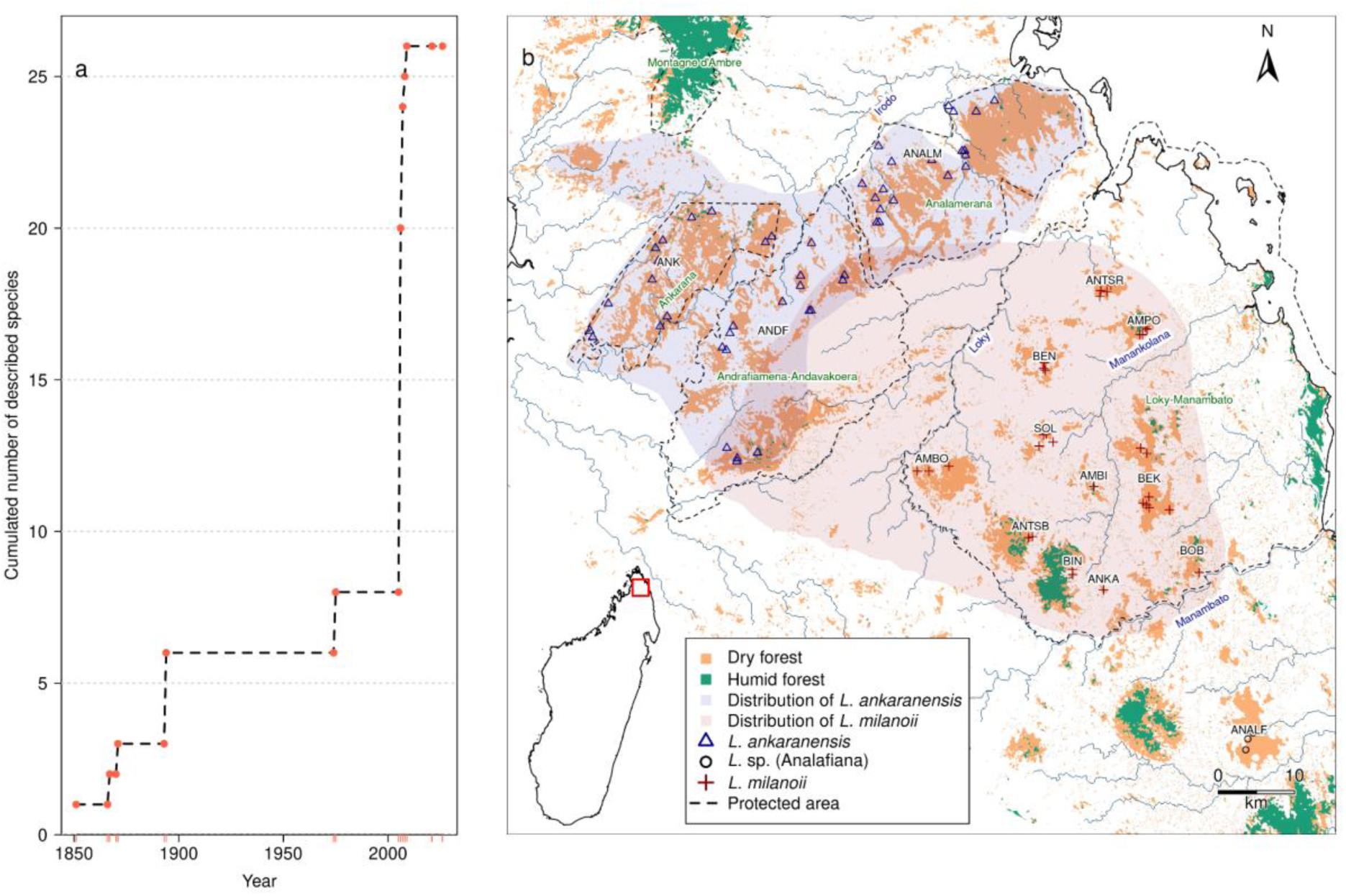
**a**) Temporal evolution of the number of described sportive lemur species (genus *Lepilemur*). This figure shows the increase in the number of species, from one to six in the 19th century, and then from eight in 2005 to 26 in 2009. **b**: Map of the described distribution ranges of *L. ankaranensis* and *L. milanoii*, of their reported area of overlap in Andrafiamena-Andavakoera, and of the samples included in the present study. Individuals in Analafiana forest (*L.* sp.) have not yet been taxonomically identified to species level. BOB: Bobankora, BEK: Bekaraoka, ANKA: Ankaramy, BIN: Binara, ANTSB: Antsahabe, AMBI: Ambilondamba, AMBO: Ambohitsitondroina, SOL: Solaniampilana, BEN: Benanofy, AMPO: Ampondrabe, ANTSR: Antaharaingy, ANDF: Andrafiamena, ANALM: Analamerana, ANK: Ankarana, ANALF: Analafiana Here, to examine the sympatric area and clarify the taxonomic status of *L. ankaranensis*, *L. milanoii* and individuals from the Analafiana forest, we use an approach inspired by the integrative taxonomy framework recently developed by van Elst et al., (2025) for the genus *Microcebus*. For that, we use genomic and morphological data from 84 *Lepilemur* individuals, encompassing the entire distribution range of two putative species, *L. ankaranensis* (n = 48) and *L. milanoii* (n = 33), with the addition of individuals from Analafiana forest (n = 3), south of the Manambato river, thought to represent the southern limit of *L. milanoii’*s distribution range (Figure 1b). Using genomic clustering, multivariate analyses, and isolation by distance comparisons, we find no evidence of a sympatric zone and show that despite the genetic differentiation between fragmented populations occurring within the range of each putative species, the genomic and morphological diversity of the *L. ankaranensis* - *L. milanoii* group is clinal, and mostly explained by geographic distance. Our results therefore suggest that *L. milanoii* is a junior synonym of *L. ankaranensis*. Furthermore, it implies that the Andrafiamena forest hosts conspecific individuals with slightly differentiated mtDNA backgrounds, rather than slightly differentiated sympatric species. On this basis we re-evaluate the conservation metrics and update the conservation status of *L. ankaranensis*, which keeps the Endangered (EN) IUCN status under the criteria B1ab(i-v).

In northern Madagascar, three species of sportive lemurs have been described: *L. septentrionalis*, *L. ankaranensis*, and *L. milanoii* (Mittermeier et al., 2023). Unexpectedly, co-occurrence of the latter two species has been reported in Andrafiamena forest, which is the central area of their common distribution range. (Figure 1b; Louis, 2006). While several cases of sympatry have been reported among Microcebus species, to our knowledge, this is the only reported case of sympatry in sportive lemurs (Mittermeier et al., 2023). Cases of sympatry among *Microcebus* species may result from secondary contact between long-diverged non-sister lineages (Kappeler et al., 2022), which most likely had time to accumulate reproductive incompatibilities (Poelstra et al., 2021; van Elst et al., 2025). Contrastingly, in the *L. ankaranensis* - *L. milanoii* case, the low mtDNA divergence suggests that little evolution has undergone between these two clades, questioning their separate taxonomic status, particularly in the context where the two clades share a significant part of their distribution area.

## Materials and methods

### Study species

*Lepilemur milanoii* and *L. ankaranensis* are two putative sportive lemur species of the family Lepilemuridae, occurring in the north of Madagascar. *L. ankanarensis* was first described by Rumpler and Albignac (1975) and subsequently confirmed by Rumpler et al. (2001) as karyotypically distinct (2N = 36-38) from *L. septentrionalis* (2N = 34-36), based on individuals sampled in Montagne d’Ambre, Ankarana, and the Andrafiamena hill chain (Figure 1b). *L. milanoii* was proposed as a new species by Louis et al. (2006) based on mtDNA data from individuals sampled in Ankarana, the Andrafiamena hill chain, and the region south of the Loky river and north of the Manambato river (the Loky-Manambato region hereafter). The mtDNA sequences formed two clades, present mainly north and south of the Loky River. The only exception to that pattern was the Andrafiamena forest, where both clades were identified and thus interpreted as a region of sympatry. The two putative species occur in dry deciduous and subhumid evergreen forests with population densities estimated in several forests and found to vary between ∼35 and ∼600 ind/km^2^ (Hawkins et al., 1990; Salmona et al., 2014). Several *Lepilemur* populations have been shown to be highly sensitive to forest fragmentation and hunting (Craul et al. 2009). Apart from their possible dependence on *Strychnos madagascariensis* tree holes as a sleeping site (Salmona et al., 2015), their ecology and behavior remain understudied.

### Study region

We visited all 17 major forests within the known distributions of the two putative *Lepilemur* species (Figure 2) during the dry season (April–November) of 2010, 2011, 2012, and 2013. Specifically, we visited the Loky-Manambato region (also known as the Daraina region), limited by the Loky and Manambato rivers (Figure 1b), the Andrafiamena-Andavakoera massif, the Ankarana National Park and the Analamerana Special Reserve. These study sites have been previously described in detail by several authors (Burivalova, 2009; Goodman & Wilmé, 2006; Hawkins et al., 1990; Ranirison, 2010). Briefly, these are mainly fragmented dry forests, frequently surrounded and/or connected by riparian corridors or smaller fragments. Yet, some forests in the southern part of the Loky-Manambato region (Binara, Antsahabe, Bobankora), as well as some valleys of Andrafiamena, and some corridors of the Ankarana Plateau harbor subhumid forests (Moat & Smith, 2007).

**Figure 2 :**
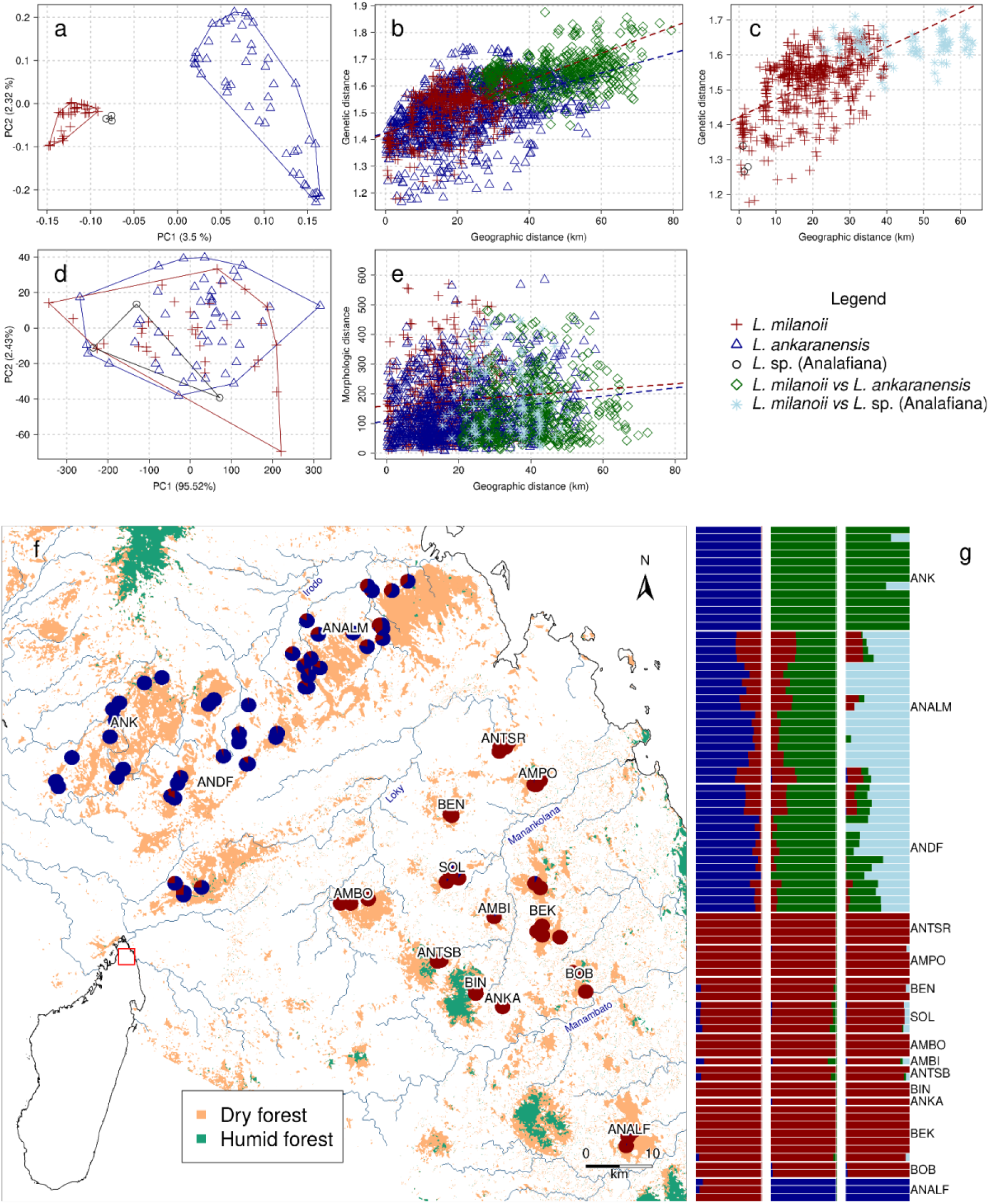
Structure patterns based on genomic and morphological data. PCA bidimensional representation of the genomic **(a)** and morphological **(d)** variability within and among candidates. Genomic **(b & c)** and morphological **(e)** patterns of isolation by distance between the candidate species *L. ankaranensis* and *L. milanoii* **(b & e)** and **(c)** *L. milanoii* and the population of the Analafiana (ANALF) forest. The within and among (*vs*) candidate species individual distances are represented with different symbols and colors, and the candidate species regressions are represented accordingly to emphasize the absence of disruption within and among candidate distances. Pie chart map **(f)** of individual genomic clustering probabilities, assuming two clusters (*K* = 2) and bar plots **(g)** displaying individual genomic clustering probabilities from K=2 to K=4. BOB: Bobankora, BEK: Bekaraoka, ANKA: Ankaramy, BIN: Binara, ANTSB: Antsahabe, AMBI: Ambilondamba, AMBO: Ambohitsitondroina, SOL: Solaniampilana, BEN: Benanofy, AMPO: Ampondrabe, ANTSR: Antaharaingy, ANDF: Andrafiamena, ANALM: Analamerana, ANK: Ankarana.

### Sample collection

We captured sportive lemurs daily, by hand, with gardening gloves in tree-hole sleeping sites, or in tree tangles using tranquilizing blowpipe syringe projectiles (Telinject, Römerberg, Germany) with ketamine hydrochloride (∼10 mg/kg). We handled sportive lemurs after tranquilizing them with an injection of ketamine hydrochloride (∼3 mg/kg). We collected sportive lemur ear biopsies from each animal and stored the samples in Queens Lysis Buffer for DNA preservation under field conditions (Seutin et al., 1991). All field handling and sampling procedures followed the Code of Best Practices for Field Primatology of the International Primatological Society and adhered to the legal requirements of France, Madagascar, and Portugal and were approved by the Malagasy Ministry of the Environment and Sustainable Development.

### DNA extraction and RAD sequencing

We extracted DNA using the DNeasy® tissue and blood extraction kit (QIAGEN©, Netherlands). We improved overnight tissue digestion by using 20 µl of 1 M DTT (DL-dithiothreitol), as described in Sgarlata et al. (2018) and Aleixo - Pais et al. (2018). We prepared restriction-site associated DNA (RAD) sequencing libraries using 40-100 ng of genomic DNA from each sample digested by the enzyme SbfI. RAD libraries were prepared following the two protocols described in (Poelstra et al., 2021; van Elst, Sgarlata, et al., 2025) detailed in Supplementary Method S2 and for each sample in Supplementary Table S2.

### Bioinformatics

Based on preliminary analyses (Salmona, 2015), we selected 84 RAD-genotyped individuals for this study, across the putative *L. ankaranensis* and *L. milanoii* distribution ranges, including the Analafiana forest, south of the Manambato river, as noted above (Figure 1b and Table S2 and S3). Raw RAD reads were demultiplexed with the “process_radtags” function of Stacks v2.2 (J. Catchen et al., 2013; J. M. Catchen et al., 2011), trimmed with Trimmomatic v0.39 (Bolger et al., 2014) (Leading: 3, Trailing: 3, Slidingwindow: 4:15, Minlen: 50), and aligned against the only assembled *L. ankaranensis* genome (GCA_963573785.1) with BWA-MEM 0.7.17 (Li & Durbin, 2009). RAD sequencing metadata and statistics are given in Supplementary Table S2 and S3. We estimated genotype likelihoods (GL) with the SAMtools model in ANGSD (Korneliussen et al., 2014). We retained (1) sites with an individual read depth larger than two, (2) sites present in at least 75% of focal individuals, (3) bases with a mapping quality larger than 20, (4) reads with a minimum mapping quality of 20, (5) biallelic variants with a probability below 1e−6 and (6) sites with a minor allele frequency (MAF) larger than 0.05.

### Species delimitation

van Elst et al., (2025) proposed an integrative taxonomy framework, which they applied to putative species of the genus *Microcebus*. This approach stresses the use of generalizable criteria based on morphological, genomic, and spatial analyses to distinguish differentiation between divergent populations from true species. Because both the *Microcebus* and *Lepilemur* genera are nocturnal primates with a similar number of species and a similarly recent increase in the number of identified species, it seems natural to apply the proposed framework to the *Lepilemur* genus. We thus used this framework to evaluate whether *L. ankaranensis* and *L. milanoii* are part of one large metapopulation in a fragmented habitat or separately evolving lineages (*i.e.,* distinct species *sensu* De Queiroz, 2007). We stress, however, that we could not apply the framework in its entirety because we lack a reference species to compare our results to. Specifically, we used an isolation by distance (IBD) approach, but we could not apply the genomic sliding-window IBD and morphological resampling IBD steps.

In brief, the retained procedure essentially relies on genomic analyses to detect population genetic structure, genetic differentiation, and gene flow among sister candidate species. We combined (i) genetic clustering and Principal Component Analysis (PCA) to evaluate if and how genetic structure fits the geography of the candidate species and suggest connectivity among clusters. We then used (ii) IBD to assess whether genetic distances between candidate species depart from a model of intraspecific spatial structure and suggest greater genetic differentiation between than within clades for the same geographic distance (Sgarlata et al., 2019). Finally, we applied (iii) PCA and IBD approaches to morphological data to similarly evaluate if it supports a model of intraspecific spatial structure. Despite some departure from the van Elst et al. (2025) procedure, we maintained their decision workflow (Figure 1 in van Elst et al., 2025), which gives priority to the genetic IBD analysis and subsequently evaluates other lines of evidence if the genetic distances between candidate species depart from a model of intraspecific spatial structure.

#### Genomic analyses

We conducted genetic PCA with PCAngsd (Meisner & Albrechtsen, 2018), and genetic clustering analyses with NgsADMIX (Skotte et al., 2013) for twenty independent runs of each value of *K* between 1 and 6. We assessed intraspecific and interspecific IBD using PCA-based genetic distance using all axes (Shirk et al., 2017) and haversine geographic distance, among individuals.

#### Morphometric analyses

We used 18 morphological variables systematically collected in the field from the 84 adult animals to assess morphometric diversity within and between *Lepilemur* species (Table S3). We controlled for potential inter-observer bias using a slightly modified version of the framework recently developed by Schüssler et al. (2024) for the genus *Microcebus*. The authors proposed a framework to reduce the inter-observer bias (i.e., increased normality of data distributions) in multi-observer measurement dataset of small morphologically cryptic mammals. Given that the variables were measured on anesthetized animals and that *Lepilemur* are one order of magnitude larger than *Microcebus*, we considered that the measurement error was lower than for *Microcebus* which were not systematically anesthetized. This led us to lower the threshold suggested by Schüssler et al. (2024) for discarding outliers down to the *Median* ± (3 ∗ *Median absolute deviation*). Furthermore, when calculating the observer coefficient of variation for each variable, observers with a low number of animals measured were not considered but were kept for subsequent analyses if not identified as outliers. We first naively summarized the major morphological variation using PCA in the R package pcaMethods (Stacklies et al., 2007). Second, we evaluated the morphological IBD, to assess whether geographic distances could explain part of the morphological variation, using PCA-based morphometric distance across all axes (Shirk et al., 2017) and haversine geographic distances, among individuals. Third, we estimated the multidimensional morphological space overlap between putative species using the *dynRB* v0.19 R package (Maislinger et al., 2015).

### Conservation status update

Based on the updated taxonomy presented here, we provide new conservation status recommendations for all valid *Lepilemur* species following the International Union for Conservation of Nature (IUCN) guidelines (IUCN, 2012). To do so, we calculated two IUCN metrics, the extent (EOO) and area of occurrence (AOO) using *terra* v.1.8-93 (Hijmans, 2020). We merged the distribution range of *L. anaranensis*, *L. milanoii* and the forest on Analafiana (Louis, Bailey, Raharivololona, et al., 2020; Louis, Bailey, Sefczek, et al., 2020) to calculate the EOO using the minimum convex polygon method. We estimated the AOO as the 2017 forest cover area encompassed by the merged distribution ranges. We further estimated *L. ankaranensis*’ potential population decline due to habitat loss (i.e., forest loss), using Madagascar’s 1953-2017 temporal forest cover reconstructions (Vieilledent et al., 2017), as a proxy for population size changes. We then compared these metrics and estimates to the official IUCN criteria (IUCN, 2012) to propose an update to *L. ankaranensis* conservation status.

## Results

The genetic analyses conducted on 84 individuals (*L. ankaranensis* n = 48, *L. milanoii* n = 33, Analafiana forest n = 3) revealed no individuals of distinct nuclear genomic background in sympatry in the Andrafiamena-Andavakoera protected area (n = 16). However, our analyses show the existence of genetic differentiation between samples collected north and south of the Loky River. The first axis of the PCA (Figure 2a) separates individuals north and south of the Loky River, although it explains less than 3.5% of the genetic variation and shows more variation within than between candidate species. A similar structure is revealed by the clustering analysis (Figure 2f,g), with distinct northern and southern clusters and most individuals from the Analamerana and Andrafiamena forests being genetically intermediate (i.e., considered “admixed” by the method). The posterior likelihood of the clustering analyses shows a near-monotonous increase from *K*=1 to *K*=10, pointing to a hierarchical decomposition of fine-scale population structure rather than interspecific distinction (Figure S1). The ΔK statistic suggests that a *K* value of two might best describe the data (Figure S1), genetically partitioning individuals between the two sides of the Loky River. For *K* ≥ 2, the clustering continues identifying genetically intermediate individuals (Figures 2f,g), which suggests past connectivity among clusters (i.e. non-null probability to belong to the same cluster) despite the sampling gap along the Loky River separating the two candidate species.

Similarly, the IBD analyses show that both the genetic variation between *L. ankaranensis* and *L. milanoi* and between *L. milanoi* and the southernmost Analafiana forest is clinal (Figure 2b,c), and substantially explained by geographic distances (*R*^2^ = 0.24 and *R*^2^ = 0.29, respectively). The IBD analyses therefore suggest that the genetic differentiation variation between the candidate species is influenced by geographic distances, and does not depart from a model of intraspecific spatial structure. Additionally, the morphometric data also supports continuous variation, with a PCA plot exhibiting almost complete overlap (overlap estimates ∼83%, Figure 2d, Table S4) of the two candidate species morphospace. The IBD analysis of morphometric data (Figure 2e) also suggests that morphometric variation between the candidate species is influenced by geographic distances and does not depart from a model of intraspecific spatial structure.

Finally, the population of the Analafiana forest (south of the Manambato River) appears genetically close to the individuals of the Loky-Manambato region (Figure 2). Furthermore, although it separates as its own cluster at *K* = 3 (Figure 2g), it does not seem to depart from a model of intraspecific spatial structure (Figure 2c) despite its apparent geographic isolation from the other populations located north of the Manambato River.

Our IBD analyses indicate that genetic distances between *L. ankaranensis* and *L. milanoii* and between *L. milanoii* and the population of Analafiana do not depart from an intraspecific model. This result alone suggests that the candidate species *L. ankaranensis*, *L. milanoii* and the population of Analafiana all belong to the same species. We, however, carefully evaluated the evidence provided by other genomic approaches, all of which suggest that although the candidate species can be distinguished from genomic data, they show signs of recent connectivity, and their differentiation is not sufficient to justify distinct species statuses. Morphometric data provide no further evidence to distinguish the candidate species. Therefore, all results suggest that *L. milanoii* is a junior synonym of *L. ankaranensis* and that the individuals of the Analafiana forest also belong to *L. ankaranensis*.

Merging the distribution range of the putative *L. ankaranensis* and *L. milanoii* species together with the population from Analafiana forest considerably expands *L. ankaranensis* extent of occurrence (EOO) from 2350 km^2^ to 4970 km^2^ and likely increases the area of occurrence (new AOO = 858.64 km²). Using forest loss between 1953 and 2017 as a proxy for population size loss, we estimate that *L. ankaranensis*’s population size likely decreased by 37.39% during that period. The EOO below 5 000 km² provides the sole but sufficient criterion, B1ab(i-v), that suggests maintaining the Endangered (EN) status of *L. ankaranensis*.

## Discussion

Our results show that the genomic and morphological differentiation of the *L. ankaranensis* -*L. milanoii* group is clinal, and mostly explained by geographic distance. The low mitochondrial genetic differences previously attributed to species divergence (Louis et al., 2006) are only partially reflected by the nuclear genomic population structure. But no individuals of distinct nuclear genomic background were found in sympatry in the Andrafiamena forest. Instead, our results confirm the existence of genetic differentiation between samples collected in the north and south of the Loky River. This population structure among the Loky River banks is in line with results from other codistributed small mammal species. For instance, *Propithecus tattersalli* and *Microcebus tavaratra* populations were found to be primarily structured by rivers and by forest fragmentation (Quéméré et al., 2010, Aleixo-Pais et al., 2018; Sgarlata et al., 2018). Similarly, major rivers were found to act as the main driver of population differentiation in *M. jonahi* and *M. lehilahitsara* ((van Elst, Schüßler, et al., 2025)). Furthermore, large rivers are major drivers of allopatric speciation in lemurs in general (e.g. Martin, 1972; Olivieri et al., 2007) and in *Lepilemur* in particular (e.g. Craul et al., 2007).

Our IBD results, however, show that none of the pairs of candidate species depart from an intraspecific model. This result clarifies that *L. milanoii* is a junior synonym of *L. ankaranensis* and that the individuals of the Analafiana forest also belong to *L. ankaranensis*. It also suggests that the Loky River is a partial barrier to gene flow in *L. ankaranensis*, in line with previous work on *Lepilemur* suggesting that large rivers do not always act as drivers of speciation (Craul et al., 2008). Furthermore, this result implies that the Andrafiamena forest, located north of the Loky River, hosts conspecific *L. ankaranensis* individuals of slightly differentiated mtDNA background, rather than individuals from two species (*L. ankaranensis* and *L. milanoii*) occurring in sympatry. This result is consistent with what we know for *Lepilemur* species across Madagascar, where no other case of species sympatry has been described yet (Mittermeier et al., 2023; Radespiel et al., 2022).

In line with previous work (e.g., van Elst et al., 2025, Poelstra et al., 2021), our results suggest that cryptic species delimitation based solely on a few mitochondrial or nuclear DNA sequences with low divergence should be considered with caution, especially when sampling is geographically limited. Nevertheless, we must stress that more data (such as genomic data) is no guarantee of accurate species delimitation. For instance, Sukumaran & Knowles (2017) showed using simulations that structured populations are often misidentified as species by popular multispecies coalescent approaches. In addition, Mason et al., (2020) showed that the latter bias was stronger when populations are sparsely sampled. Explicitly considering geography has repeatedly been suggested as a way to overcome this problem (e.g., Hausdorf, 2025; Sgarlata et al., 2024; van Elst, Sgarlata, et al., 2025), and we advocate its systematic integration in discussions about the identification of new species, particularly when the number of sampled localities is limited.

In the genus *Lepilemur*, interestingly, before the 2006-2009 taxonomic burst, most species were described based on either morphological (e.g., Jungers & Rumpler, 1976) or karyotypic differences (e.g., Rumpler et al., 2002). Our results suggest that a careful and geographically informed examination of genomic data should be conducted throughout the island and may result in a decrease in the number of *Lepilemur* species, similar to what was recently suggested for mouse lemurs (genus *Microcebus*, van Elst et al., 2025). For instance, *L. mittermeieri* was synonymized with *L. dorsalis* by Roos et al., (2007) but kept as distinct species by both the reference field guide (Mittermeier et al., 2023) and the IUCN Red List. Considering the similarity of the mtDNA material found in both species and their geographic proximity, *L. mittermeieri* could be confirmed as a junior synonym when population genomic data become available.

At the same time, we must stress that each genus may have rather different evolutionary histories, and we should remain open to the fact that the number of *Lepilemur* species may change in unexpected directions. Indeed, previous karyotypic studies have shown that species of the genus *Lepilemur* exhibit the most differentiated karyotypes of all lemur genera, with significant within-species variation (Rumpler et al., 2008). Much work is thus needed to clarify and document the full genomic diversity of sportive lemurs and to identify the drivers of their diversification.

From a conservation perspective, synonymizing *L. milanoii* with *L. ankaranensis* and attributing the population of Analafiana to *L. ankaranensis* considerably extends its distribution range. Although the conservation status of *L. ankaranensis* remains “Endangered” (EN) under the criteria B1ab(i-v), the updated taxonomy may change the conservation narrative used by the managers of the Loky-Manambato protected area (PA, the NGO Fanamby). To that purpose, we defined three main evolutionary significant units (Andrafiamena-Andavakoera-Ankarana region, Loky-Manambato region, and Analafiana forest) that can be leveraged to emphasize the specificity of each particular area, although the *Lepilemur* populations of the Loky-Manambato belong to *L. ankaranensis* and are no longer endemic to the PA.

## Supplementary information

Supplementary figures:

https://docs.google.com/document/d/1RLp-PtWlfbeGFlFZbk-xNmdfO_KDmbkBd7WdxsNAS3Q/edit?usp=sharing

Supplementary tables:

https://docs.google.com/spreadsheets/d/1VHBMSwzdy3DqlC16ghp0ruTXZQ98m0cTQ6AcF7Yiqhg/edit?usp=sharing

## Data Sharing and Accessibility

All data, scripts and intermediary files necessary to replicate the analyses have been (will be) deposited in the European Nucleotide Archive (https://www.ebi.ac.uk/ena/browser/home, SRAXXXXXX), Github (https://github.com/besoa/ms_Lepilemur_ankaranensis_2026), Zenodo (link), respectively.

## Acknowledgements

We thank CAFF/CORE, the Direction Générale de la Gouvernance Environnementale (DGGE), Madagascar National Parks, the Fanamby NGO (including S. Rajaobelina and V. Rasoloarison), the Direction Régionale de l’Environnement et du Développement Durable (DREDD) of DIANA and SAVA administrative regions. The fieldwork was possible thanks to the continuous support of the Département de Biologie Animale et Ecologie, University of Mahajanga, and the University of Antsiranana. We also thank the many exceptional local guides and cooks for their help in the field and for sharing their incomparable expertise of the forest. Financial support for this study was provided by the ‘Institut de Recherche pour le Développement’ (IRD-ARTS to MBRR and ER) ‘Fundação para a Ciência e a Tecnologia’ (ref. SFRH/BD/64875/2009 to JS and PTDC/BIA-BIC/4476/2012 PTDC/BIA-BEC/100176/2008 to LC), the GDRI Madagascar, the Laboratoire d’Excellence (LABEX) entitled TULIP (ANR-1 0-LABX-41), the Rufford Small Grant Foundation, grant 10941-1 to JS and 12973-1 to MTI, the Instituto Gulbenkian de Ciência, the IRP BEEG-B (International Research Project - Bioinformatics, Ecology, Evolution, Genomics and Behaviour) (CNRS) to LC and Optimus!Alive-IGC fellowship to CKR and HT. We are grateful to the Genotoul bioinformatics platform Toulouse Occitanie (Bioinfo Genotoul, https://doi.org/10.15454/1.5572369328961167E12) for providing computing and storage resources. This study was conducted in agreement with the laws of the countries of Portugal, France, and Madagascar.

