## Supplementary Figures for "Exploring the only known case of sympatry in sportive lemurs: isolation by distance or speciation?"

**<u>Supplementary material for</u>** Salmona, Ranjavao et al., in prep. Exploring the only known case of sympatry in sportive lemurs: isolation by distance or speciation?

### <u>Authors and affiliations:</u>

### <u>Table of content:</u>

**S1: Clustering likelihood and ∆K dynamics** **1**
**S2: Library preparation** **2**

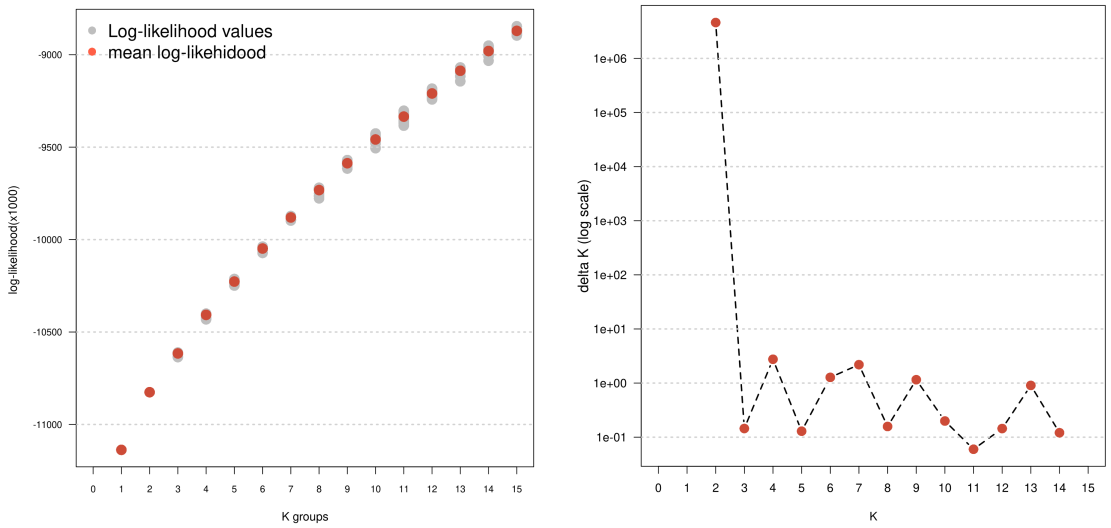


### <u>S1: Clustering likelihood and ∆K dynamics</u>

Posterior likelihood (left panel) and deltaK (right panel) dynamics for increasing number of clusters (*K*). The posterior likelihood of the clustering analyses conducted with ngsAdmix, shows a near-monotonous increase from *K*=1 to *K*=10, pointing to a hierarchical decomposition of fine-scale population structure rather than interspecific distinction. The ∆K statistic suggests that a *K* value of two might best describe the data.

### S2: Library preparation

RAD libraries were prepared following the two protocols described in Poelstra et al. 2021 and van Elst et al., 2025 and the detail for each sample is reported in Table S2.

**1. Oregon:** Library preparation was based on Genomic resources development consortium et al., (2015). Specifically, 40–100 ng of extracted genomic DNA were digested with the SbfI restriction enzyme (New England Biolabs) and subsequently ligated to the P1 adapters 73 . Up to 48 samples were pooled into sub-libraries and sheared for 5 min using a Bioruptor for 6 min to an average target size of 500 bp. Next, end-repair and 3' adenylation were performed, P2 adapters were ligated, and libraries were amplified in 14 cycles of PCR. Finally, sub-libraries were purified with AMPure XP beads (Agencourt), pooled based on yield, and single-end sequenced (100 bp, 48 individuals/lane) on an Illumina HiSeq 2000 at the University of Oregon Core Facility.

**2. Toulouse:** Library preparation was also based on Genomic resources development consortium et al., (2015) In contrast to the protocol mentioned above, 40–200 ng of genomic DNA were used, sub-libraries were sheared for 45 s in Covaris® M220, only 10 PCR cycles were conducted, and sequencing was performed on an Illumina HiSeq 3000 (paired-end, 150 bp, 96 individuals/lane) at the GenoToul Sequencing Platform Facility (Toulouse, France).
